# Profiling the LAM family of contact site tethers provides insights into their regulation and function

**DOI:** 10.1101/2024.04.18.590074

**Authors:** Emma J. Fenech, Meital Kupervaser, Angela Boshnakovska, Shani Ravid, Inês Gomes Castro, Yeynit Asraf, Sylvie Callegari, Christof Lens, Peter Rehling, Maya Schuldiner

## Abstract

Membrane contact sites are molecular bridges between organelles that are sustained by tethering proteins and enable organelle communication. The endoplasmic reticulum (ER) membrane harbors many distinct families of tether proteins that enable the formation of contacts with all other organelles. One such example is the LAM (<u>L</u>ipid transfer protein <u>A</u>t <u>M</u>embrane contact sites) family, composed of six members, each containing a lipid binding and transfer domain and an ER-embedded transmembrane segment. The family is divided into three homologous pairs each unique in their molecular architecture and localization to different ER subdomains. However, what determines the distinct localization of the different LAMs and which specific roles they carry out in each contact are still open questions. To address these, we utilized a labeling approach to profile the proximal protein landscape of the entire family. Focusing on unique interactors we could support that Lam5 resides at the ER-mitochondria contact site and demonstrate a role for it in sustaining mitochondrial activity. Capturing shared interactors of multiple LAMs, we show how the Lam1/3 and Lam2/4 paralogous pairs could be associated specifically with the plasma membrane. Overall, our work provides new insights into the regulation and function of the LAM family members. More globally it demonstrates how proximity labeling can help identify the shared or unique functions of paralogous proteins.

## Introduction

Eukaryotic complexity is afforded by cell compartmentalization into membrane-bound organelles, each of which is defined by a unique protein and lipid landscape organized into specific substructures. However, these organelles must communicate with one another to coordinate cellular processes and output. This can be achieved through membrane contact sites – areas of close apposition (typically ∼30nm or less) between specialized membrane subdomains of different organelles, which are tethered to each other via protein-protein or protein-lipid interactions (Scorrano et al., 2019).

All organelles form contacts with each other (Shai et al., 2018) and this has been well documented for the endoplasmic reticulum (ER), which forms multiple and expansive contacts with different organellar membranes (Valm et al., 2017). Lipid transfer is an indispensable function of ER contact sites since the majority of lipid biosynthetic machinery is resident in the ER membrane. One protein domain capable of mediating the transfer of the essential lipid, sterol, is the <u>St</u>eroidogenic <u>A</u>cute <u>R</u>egulatory <u>T</u>ransfer (StART) domain. In *Saccharomyces cerevisiae* (called yeast from here on) there is a family of StART domain proteins that resides in the ER and is composed of three homologous pairs: Ysp1 (Lam1) and Sip3 (Lam3); Ysp2 (Ltc4 or Lam2) and Lam4 (Ltc3); and Lam5 (Ltc2) and Lam6 (Ltc1) (Figure 1A) (Gatta et al., 2015). For simplicity and consistency, we will henceforth refer to these proteins only by their LAM nomenclature. Apart from their StART-like domains, these proteins also share an N-terminal <u>P</u>leckstrin <u>H</u>omology (PH) domain, and a C-terminal transmembrane domain (TMD) which anchors them in the ER membrane (Gatta et al., 2015). The presence of these domains is conserved to the family of human ASTER proteins; ASTER-A, ASTER-B and ASTER-C, which are also called GRAMD1A, GRAMD1B and GRAMD1C (Gatta et al., 2015; Sandhu et al., 2018).

**Figure 1:**
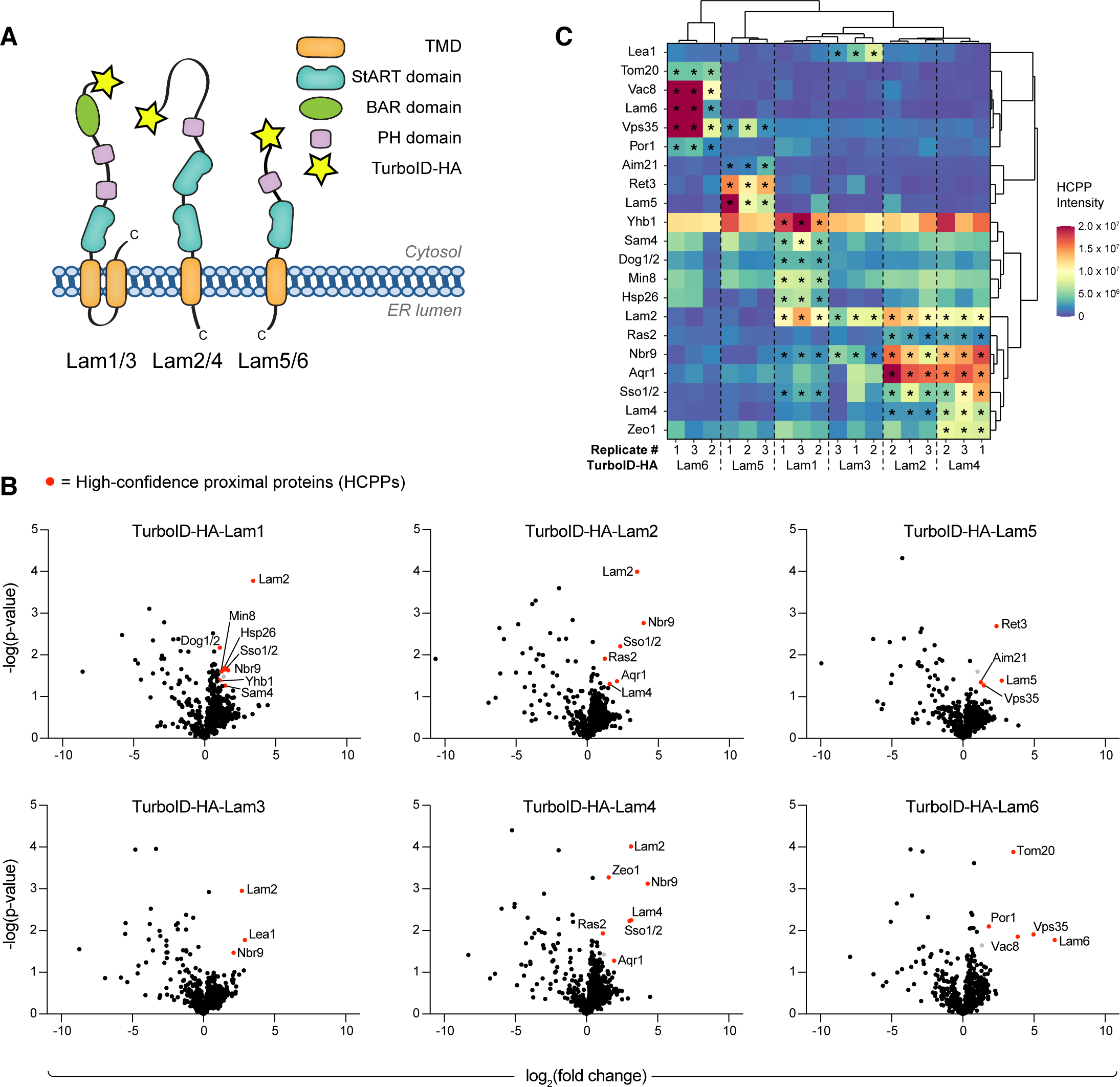
Proximity profiles of LAM family members differentiate between shared and unique features. (A) Schematic of the yeast LAM family, shown as three pairs of homologs: Lam1 (Ysp1) and Lam3 (Sip3); Lam2 (Ysp2/Ltc4) and Lam4 (Ltc3); Lam5 (Ltc2) and Lam6 (Ltc1). Highlighted are: the cytosolic-facing TurboID-HA tags (yellow) at the N-termini; the BAR (<u>B</u>in-<u>A</u>mphiphysin-<u>R</u>vs; green), PH (purple) and StART (blue) domains; and the C-terminal TMDs (orange). (B) Volcano plots showing -log(*P-*value) against log_2_(fold-change) for all proteins identified by LC-MS/MS of the TurboID-HA-tagged samples. High-confidence proximal proteins (HCPPs) enriched in the appropriate TurboID-HA-LAM samples are marked with their names and highlighted in red; proteins which passed the criteria for *P*-value and fold-change, but were only identified by one unique peptide, are colored grey. (C) Hierarchical clustering of the TurboID-ID LAM biological triplicates and their HCPPs, shown as a heatmap. The colors are defined by the normalized intensity. The HCPPs highlighted in red in (B) are marked by an asterisk. The data shows clustering according to the biological replicates. The homologous LAM pairs cluster together, with Lam1-4 separate from Lam5 and Lam6.

The LAM proteins were discovered to localize at distinct ER contact sites. Lam1,2,3 and 4 (Lam1-4) are present at ER-plasma membrane (PM) contacts (Gatta et al., 2015; Murley et al., 2017), Lam5 at ER-mitochondria (Gatta et al., 2015) and ER-Golgi contacts (Weill et al., 2018a), and Lam6 at contacts between ER-mitochondria, and nuclear-vacuole junctions (NVJs) which are contacts between the nuclear ER and the vacuole (Elbaz-Alon et al., 2015; Gatta et al., 2015; Murley et al., 2015). Numerous structure-function studies on the Lam2 and Lam4 StART-like domains have mechanistically defined sterol binding, solubilization and retrograde transfer from the PM to the ER (Gatta et al., 2015; Gatta et al., 2018; Horenkamp et al., 2018; Jentsch et al., 2018; Khelashvili et al., 2019; Tong et al., 2018). However, the best characterized member of this family is Lam6, and its presence at multiple membrane contact sites makes it essential for their regulation and cross-talk (Elbaz-Alon et al., 2015). Furthermore, as opposed to Lam1-4, Lam6 mediates anterograde sterol transfer (away from the ER) required for stress-induced formation of a vacuolar membrane subdomain (Murley et al., 2015) by forming a tethering interaction with the vacuolar protein, Vac8. At the ER-mitochondria contact, Lam6 tethers both organelles by interacting with the mitochondrial protein, Tom70 (Elbaz-Alon et al., 2015; Murley et al., 2015).

The fact that Lam6 requires protein tethers on the adjacent membranes of either mitochondria (Tom70) or the vacuole (Vac8) is in some ways curious, given the presence of its N-terminal PH-like/GRAM domain. These domains are classically associated with binding anionic lipids such as phosphatidylserine (PS) and different phosphoinositide (PI) species in the adjacent membrane of the contact (Ercan et al., 2021; Naito et al., 2019; Sandhu et al., 2018). However, analysis of the LAM family PH domains shows that they expose fewer basic residues relative to typical PH domains (Tong et al., 2018). This would result in a lower affinity to anionic lipids and might underly their requirement for proteinaceous tethering molecules.

Short of Lam6, there is much less information on how other LAMs bind adjacent membranes. Two proteins, Laf1 and Dgr2, were identified as interactors of Lam1-4 (Murley et al., 2015; Topolska et al., 2020). However neither of these are integral membrane proteins and their deletion did not affect the localization of Lam2 or Lam4 at ER-PM contacts (Topolska et al., 2020). Further still, Lam5 has not been well characterized, neither in terms of interactors, nor function. Therefore, to better understand the functional significance of the LAM proteins, and how their positioning at specific ER subdomains is determined, we turned to our recently-developed enhanced proximity-labeling approach (Fenech et al., 2023) to uncover proximally residing proteins. Through this approach, we were able to shed light on a bioenergetic role for the lesser-characterized Lam5 protein, and also identify a positive regulator of Lam1-4-based ER-PM tethering, which we propose reconciles protein-and lipid-mediated binding to membranes.

## Results

### Proximity profiles of LAM family members differentiate between shared and unique features

The family of yeast LAM proteins act as contact site tethers. LAMs are anchored to the ER membrane by C-terminal TMDs and have long N-terminal domains that extend away from the ER to make contact with different membranes (Gatta et al., 2015). How these ER-resident proteins are directed to different contacts, and which cellular functions they support at these locales, is still not fully understood. We hypothesized that clues to this could come from mapping their precise protein environment at these points of contact.

To map their interactome, we turned to the efficient proximity-labeling enzyme, TurboID (Branon et al., 2018). TurboID conjugates biotin onto available lysine (K) residues in proteins within ∼10nm, allowing them to be effectively captured by streptavidin and later detected by mass spectrometry (MS). Therefore, we utilized strains expressing TurboID-HA N-terminal fusions for each of the six LAM family members (Figure 1A, Supplementary figure 1) that were part of the whole-proteome yeast TurboID library (Fenech et al., 2023). Our reasoning was that N-terminal fusions would be poised to label putative tethering machinery on the adjacent contact site membranes. Furthermore, the strains were constructed on the background of the ABOLISH (<u>A</u>uxin-induced <u>BiO</u>tin <u>LI</u>gase dimini<u>SHi</u>ng) system, which enhances the detection of TurboID-labeled protein interactors by specifically reducing levels of endogenously biotinylated proteins (Fenech et al., 2023).

These strains, together with a control strain expressing TurboID-HA-Emc6 (an ER-resident protein not associated with the LAM family), were subject to streptavidin affinity purification (AP) and analyzed by MS. To determine high-confidence proximal proteins (HCPPs) for each of the LAMs, their MS profiles were compared to that of the Emc6 sample and enriched proteins were determined as having: a *P*-value ≤ 0.05; a fold-change ζ 2; and at least two unique peptides (Figure 1B, Supplementary table 1).

The first striking observation was the strong overlap between the HCPPs of Lam1-4, driving the clustering of these four LAMs together (Figure 1C). This was not surprising since these proteins are all known to mediate ER-PM contacts (Gatta et al., 2015; Murley et al., 2017). Furthermore, the paralog pairs Lam1/3 and Lam2/4 formed their own subclusters, with a very high degree of overlap shared between the Lam2 and Lam4 HCPPs (Figure 1B, 1C). While Lam1 and Lam3 did share HCPPs, Lam1 enriched more unique soluble co-factors (Hsp26, Yhb1, Dog1/2 and Sam4) relative to Lam3 (and indeed relative to Lam2 and Lam4). Promisingly, many shared HCPPs of Lam1-4 are known to be PM-localized (Sso1/2, Nbr9, Ras2, and Aqr1) and Lam2 was identified as an HCPP of Lam1, Lam3 and Lam4; the latter being reciprocally found as a Lam2 HCPP (Figure 1B, 1C). The interconnectivity between these LAM family members has been previously reported in independent proteomic experiments performed on C-terminally tagged LAM proteins (Murley et al., 2017). Furthermore, this feature extends to the human GRAMD orthologs (Naito et al., 2019) and we could also reproduce it using FLAG-tagged constructs of GRAMD1A (Supplementary table 2) (Naito et al., 2019).

The Lam5/6 paralog pair form their own separate clusters (Figure 1C). As expected, mitochondrial proteins were enriched in the Lam6 sample, including: the outer membrane voltage-dependent anion channel, Por1; and Tom20, a component of the mitochondrial TOM (translocation of outer membrane) complex. Por1 was previously isolated together with Lam6 (Murley et al., 2015), whereas Tom20 forms a complex together with Tom70; the known tethering partner for Lam6 at ER-mitochondria contact sites. Interestingly, Lam5 was also associated with HCPPs linked to different aspects of mitochondrial biology and these are discussed below. Lastly, the vacuolar protein, Vac8, and the ζ-subunit of the coatomer complex, Ret3, were identified as unique HCPPs of Lam6 and Lam5, respectively (Figure 1B, 1C). This is consistent with their known roles: Lam6-Vac8 is a known tethering pair at the ER-vacuole contact site (Elbaz-Alon et al., 2015; Murley et al., 2015); and Lam5 localizes to the ER-Golgi interface (Weill et al., 2018a), where the coatomer machinery resides. GRAMD1A-FLAG analysis also recovered a coatomer component (Supplementary table 2), increasing our confidence in this putative interaction.

Altogether, our TurboID/ABOLISH approach uncovered known LAM family interactors, as well as many novel putative interactors that could provide clues to the function, regulation and tethering properties of these contact site proteins.

### A new role for Lam5 in mitochondrial activity at the ER-mitochondria contact site

While Lam6 is known to play a role in contact site cross-talk (Elbaz-Alon et al., 2015) and

vacuole membrane domain formation (Murley et al., 2015), its close paralog, Lam5, remains uncharacterized in terms of function. The observation that stress-induced domain formation in the vacuolar membrane is perturbed in a *Δlam6* strain where Lam5 is still present (Murley et al., 2015) suggests that their roles are at least partially non-redundant. Indeed, the HCPPs of these two LAM family members were largely unique, with only Vps35 being shared (Figure 2A). To begin exploring any overlapping and/or unique functions of Lam5 and Lam6, we first visualized GFP-tagged versions of these proteins together with vacuolar and mitochondrial markers (Figure 2B). It was immediately clear that while both proteins displayed a punctate pattern, Lam5, but not Lam6, also showed a more typical ER signature (defined as having a ‘ring’ around the nucleus and another around the cell periphery). Furthermore, there was a high degree of colocalization between the signal from the stained organelles and GFP-Lam6, as expected (Elbaz-Alon et al., 2015; Gatta et al., 2015; Murley et al., 2015). We did, however, find some colocalization of GFP-Lam5 with both vacuole and mitochondria, the latter supporting the previously observed colocalization between Lam5 and Tom6 (Gatta et al., 2015) and recent data analyzing the ER-mitochondria contact (Fujimoto et al., 2023).

**Figure 2.**
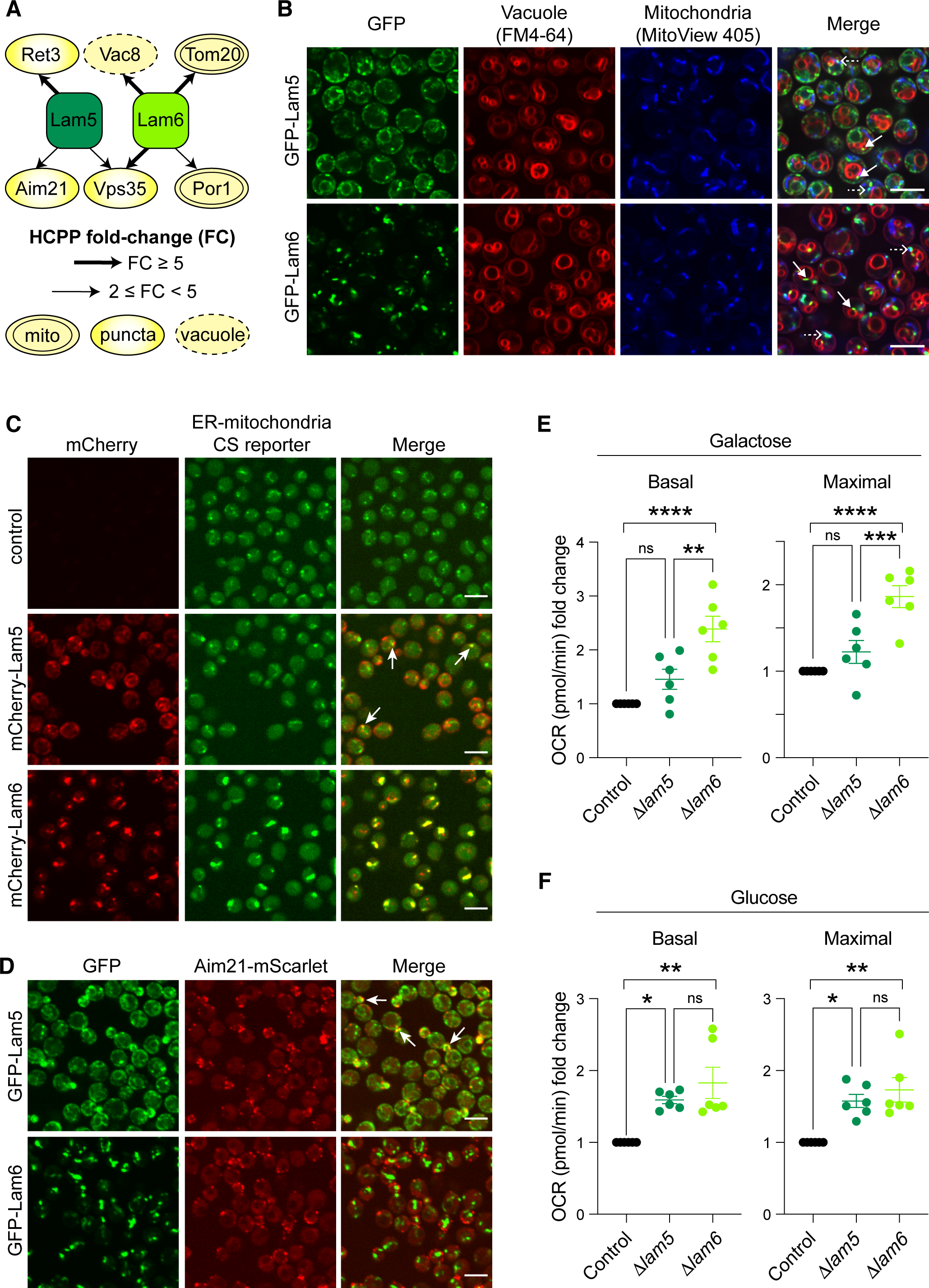
Interactome analysis reveals a new role for Lam5 in the ER-mitochondria contact site. (A) An illustration of unique and shared HCPPs of Lam5 and Lam6. HCPP fold-change is marked by a thick (fold-change ≥ 5) or thin (2 ≤ fold-change < 5) arrow, and HCPP localization/pattern (mitochondria – mito; vacuole; or puncta) is annotated according to images of the GFP SWAT libraries (Meurer et al., 2018; Weill et al., 2018b; Yofe et al., 2016) collected on Yeast RGB (Dubreuil et al., 2018). (B) Enhanced resolution confocal microscopy images of strains expressing Lam5 or Lam6 tagged on their N-termini with GFP, stained with vacuolar (FM™4-64) and mitochondrial (MitoView™405) dyes. While it was known that Lam6 localizes to contact sites with both mitochondria and vacuoles, we could also visualize Lam5 colocalization with both the vacuole and mitochondria, as highlighted by solid and dashed arrows, respectively. Scale bars are 5μm. (C) Confocal microscopy images of strains expressing Lam5 or Lam6 tagged on their N-termini with mCherry and under expression of the strong *TEF2* promoter, together with a split-Venus reporter for ER-mitochondria contact sites. Overexpressed Lam6 colocalizes with the reporter and even enhances the extent of the contact. Overexpressed Lam5 partially colocalizes with the reporter, as indicated by the white arrows. Scale bars are 5μm. (D) Confocal microscopy images of strains expressing Lam5 or Lam6 tagged on their N-termini with GFP, together with Aim21, that was identified as a putative Lam5 interactor, tagged on its C-terminus with mScarlet. Indeed, Lam5, but not Lam6, colocalizes with Aim21, especially in/around the bud, as indicated by white arrows. Scale bars are 5μm. (E) Graphs showing the fold-change in basal and maximal oxygen consumption rate (OCR) of *Δlam5* (dark green) and *Δlam6* (light green) strains relative to control (black), grown in the non-fermentable carbon source, galactose, measured by Seahorse assay (Agilent). Only the *lam6* knockout shows a significant increase in both basal and maximal OCR. The data are from six biological replicates and significance was calculated using two-way ANOVA with Tukey’s multiple comparison where: * is *p*≤0.05; ** is *p*≤0.01; *** is *p*≤0.001; **** is *p*≤0.0001; and ns is not significant. Error bars show the standard error of the mean (SEM). (F) As in (E) however strains were grown in glucose, where now both knockout strains show significantly elevated basal and maximal OCR.

To examine whether Lam5 could be a *bona fide* resident of the ER-mitochondria contact site, we assessed its colocalization with an ER-mitochondria contact site reporter based on the split-Venus probe (Shai et al., 2018). Indeed, overexpressed mCherry-Lam5 signal colocalized with the reporter at discrete puncta (Figure 2C). Lam6 tagged with mCherry also colocalized with the ER-mitochondria contact site reporter and its over-expression increased the area of the reporter’s signal, a phenomenon associated with some tethering proteins (Eisenberg-Bord et al., 2016). While Lam6 forms a contact site tether with components of the TOM complex (Elbaz-Alon et al., 2015; Murley et al., 2015) we did not identify mitochondrial outer membrane proteins as Lam5 HCPPs (Figure 1B, 1C, 2A). We did, however, identify Aim21, an actin-associated protein needed for correct mitochondrial migration and inheritance (Hess et al., 2009; Shin et al., 2018). Strikingly, GFP-Lam5 strongly colocalized with Aim21-mScarlet in the bud and bud neck region, unlike GFP-Lam6 which showed no colocalization with this protein (Figure 2D).

Our observations suggested that Lam5, in addition to Lam6, would have a function at ER-mitochondria contacts. If so, its loss should affect mitochondrial activity. To measure this, we used real-time respirometry to measure the oxygen consumption rate (OCR) in cells lacking either Lam5 or Lam6 and compared them to control cells. When grown on a non-fermentable carbon source (galactose), only the loss of Lam6 significantly affected respiration (Figure 2E, Supplementary figure 2A). However, when grown on glucose as the carbon source (the condition in which we performed our interactome profiling and microscopic analyses) we observed that both *Δlam5* and *Δlam6* had a higher basal and maximal OCR. This demonstrates that these mutations cause elevated levels of respiration and electron transport chain (ETC) activity (Figure 2F, Supplementary figure 2A). Our data uncover a previously unappreciated role for Lam5 at the ER-mitochondria contact site and suggest that the Lam5/6 paralogs differentially influence mitochondrial function through unique interactions.

### Proteins proximal to Lam1-4 reveal a novel mode of association to the PM

In addition to the Lam5/6 paralogs, there are two other homologous protein pairs; Lam1/3 and Lam2/4, which reside at ER-PM contacts (Gatta et al., 2015; Murley et al., 2017). As tethers they should have binding partners on the PM, yet their identity to date has not been uncovered. Encouragingly, we observed an enrichment of PM-resident proteins as HCPPs (Figure 1B, 1C). Furthermore, it seemed that the Lam1/3 paralog pair was less strongly associated with these proteins relative to the Lam2/4 paralogs (Figure 3A). This aligned closely with what was observed by imaging. GFP-tagged Lam1/3 display a more typical ER pattern, with both perinuclear and cell peripheral domains visible (Figure 3B), as opposed to GFP-Lam2/4, which appear exclusively as peripheral ER, indicating a tighter association to the PM. Imaging of both mitochondria and vacuoles showed essentially no overlap with these members of the LAM family (Supplementary figure 3A), highlighting the non-redundancy between Lam1-4 and Lam5/6.

**Figure 3.**
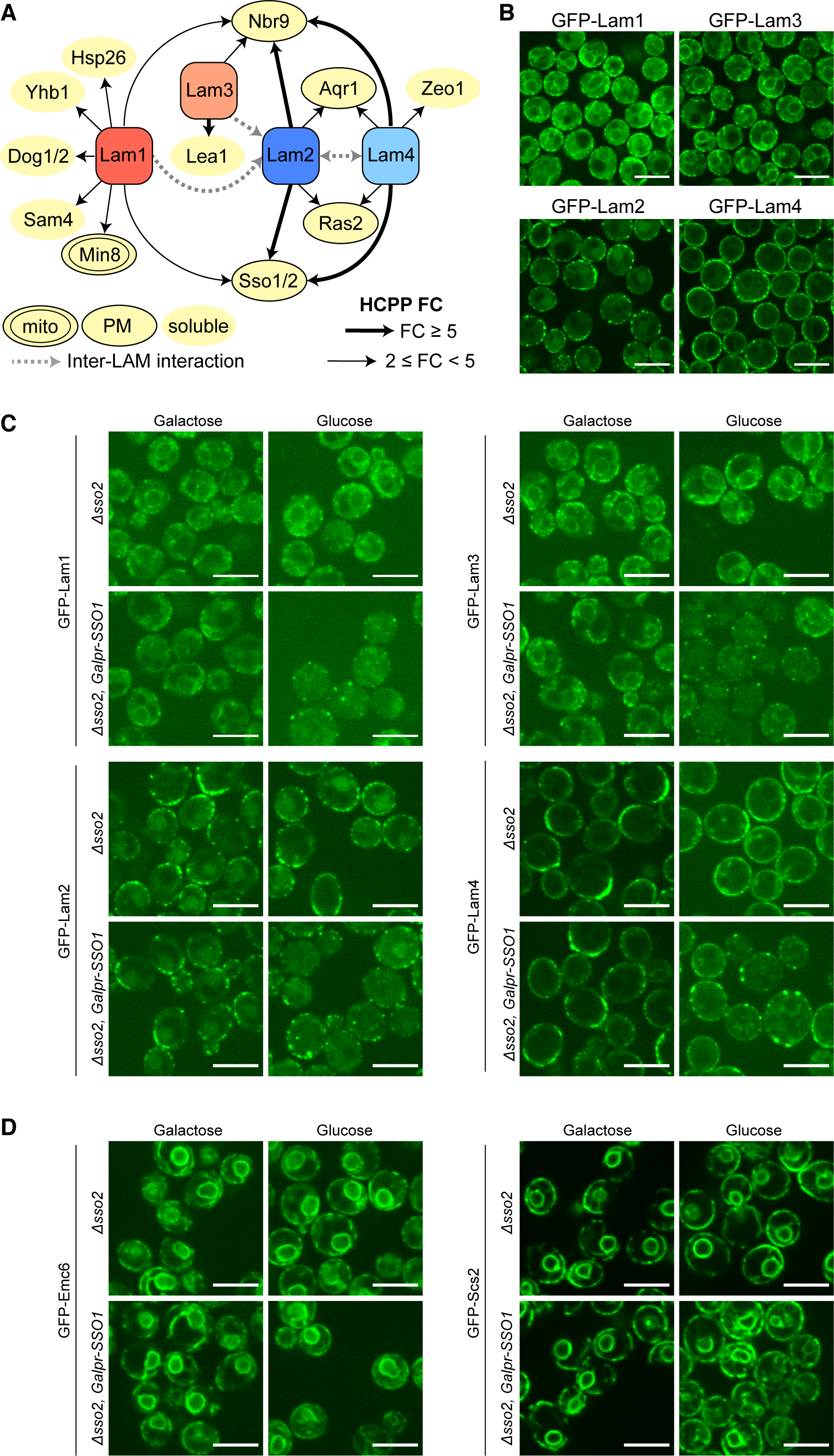
Proteins proximal to Lam1-4 reveal a novel mode of association to the PM. (A) Illustration of unique and shared HCPPs of Lam1-4. HCPP fold-change is marked by a thick (fold-change ≥ 5) or thin (2 ≤ fold-change < 5) arrow, with interactions between different LAM proteins marked by the grey, dashed line. HCPP localization (plasma membrane - PM; cyto-/nucleo-plasmic – soluble) is annotated according to images of the GFP SWAT libraries (Meurer et al., 2018; Weill et al., 2018b; Yofe et al., 2016) collected on Yeast RGB (Dubreuil et al., 2018). (B) Enhanced resolution confocal microscopy images of strains expressing Lam1-4 tagged on their N-termini with GFP. All display at least some peripheral localization (as reported) with the paralog pair Lam2 and Lam4 situated exclusively at the periphery relative to the Lam1 and Lam3 paralog pair. Scale bars are 5μm. (C) Enhanced resolution confocal microscopy images of strains grown in either galactose or glucose, expressing Lam1 and Lam3 (top panels), and Lam2 and Lam4 (bottom panels) tagged on their N termini with GFP, on the background of either *Δsso2* or *Δsso2* with an inducible/repressible promoter (*GALpr*) driving the expression of its homolog *SSO1* (*Δsso2* + *GALpr-SSO1*). In glucose, where *SSO1* is repressed creating a double mutant of *SSO1/2*, each of the tested LAM proteins loses their normal peripheral ER pattern and instead are now localized to a few bright puncta around the periphery. Scale bars are 5μm. (D) As in (C) however GFP-tagged proteins are Emc6 and Scs2 as negative controls. In glucose, the peripheral ER localization of both persists. Scale bars are 5μm.

Of the potential PM-resident binding proteins, we focused on the SSOs (paralogs Sso1 and Sso2, orthologous to human Syntaxin1A, STX1A). These proteins were found as HCPPs of Lam1-4 and have already been implicated in ER-PM contact site formation in both yeast and humans (Petkovic et al., 2014). Importantly, none of the other known ER-PM tethers (Filseck et al., 2015; Manford et al., 2012), including the SSO proteins, had been physically associated to the LAM proteins before. Since the SSOs must be in areas of close apposition between the ER and PM, we hypothesized that they may play a role in the Lam1-4-mediated ER-PM contacts. To test this, we generated *SSO* mutant strains on the background of the GFP-tagged Lam1-4. Since the double knockout of *SSO1/2* is lethal, we created either *Δsso2* strains or *Δsso2* strains which also harbored *SSO1* under regulation of a galactose-inducible/glucose-repressible promoter (*GAL1pr-SSO1*). When the latter strain is cultured in glucose-containing media, *SSO1* expression is shut-off, enabling the generation of cells with minimal levels of SSO proteins. We found that only when the *Δsso2*/*GAL1pr-SSO1* cells were grown in glucose (Figure 3C, bottom right image in each panel), did the localization of GFP-Lam1-4 proteins change dramatically; supporting the hypothesis that the SSO proteins play a role in tethering of Lam1-4 at the PM. Here, the majority of the signal, attributed to the ER-PM contact, disappeared, leaving only very few small puncta which were mainly situated around the cell periphery.

To control for non-specific effects on either the ER membrane as a whole, or the general ER-PM contact site structure, we imaged either a GFP-tagged ER membrane protein *not* involved in contact site formation (GFP-Emc6) or the yeast ortholog of VAMP-associated protein (VAP); an ER-PM contact site protein not associated with the LAM family members (GFP-Scs2). These tagged proteins, which were imaged on the same genetic background as the GFP-Lam1-4 strains, were not affected by SSO protein loss (Figure 3D). Taken together, these data suggest that SSO proteins are important, either directly or indirectly, for Lam1-4 binding the PM at ER-PM contact sites (Figure 4).

**Figure 4.**
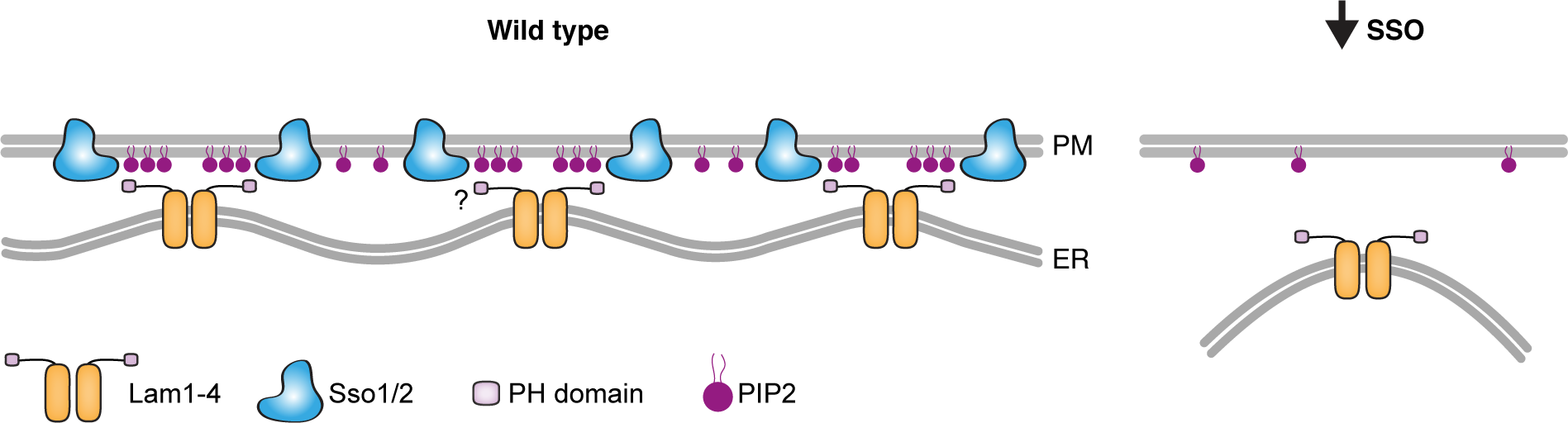
A model of how SSO proteins may affect Lam1-4 association at the ER-PM contact. A hypothesis for how SSO loss could affect LAM association with the PM: ER localized Lam1-4 may oligomerize, potentially through Lam2, bringing together their N-terminal PH and StART domains (for simplicity, only the PH domain is depicted). The PH domain can interact with PIP2 on the PM, which has been shown to be corralled by the human SSO orthologs (Honigmann et al., 2013). These local concentrations of PIP2, potentially together with Lam1-4-SSO protein-interaction (shown as a question mark), may act to increase the avidity of binding between Lam1-4 and the PM. On the contrary, when SSO protein levels are limiting, PIP2 is randomly diffused within the PM, reducing its effective concentration at contact sites and decreasing the binding capacity for the LAMs, hence reducing the amount of contacts that they form.

## Discussion

Using the newly-developed TurboID/ABOLISH system (Fenech et al., 2023) we set out to uncover novel proximal proteins for the entire ER-resident LAM family of contact site proteins, to gain a deeper understanding of their unique and shared features, and how these define their localization or regulation. Since the LAM proteins are anchored in the ER membrane by C-terminal TMDs, and their N-termini extend towards adjacent membranes to form a contact, we tagged these proteins N-terminally (Figure 1A) to capture putative interactors which may be involved in membrane tethering. While proximity-labeling methods are distinct from classical immunoprecipitation (IP), we could still recapitulate previously identified interactors including the Lam6-Vac8 interaction (Elbaz-Alon et al., 2015; Murley et al., 2015) and the association between ER-PM LAM members (Murley et al., 2017). As well as verifying known interactions, we also identified new putative proximal proteins, which we explored further.

From the proximity profiling, it was evident that the paralog pair Lam5 and Lam6 were the most distinct – both from the rest of the LAM family, and from each other (Figure 1C). Nevertheless, inspection of their localization revealed some overlap in terms of the organelles they are adjacent to (Figure 2B). Intriguingly, Lam5 showed partial colocalization with mitochondria, and encouragingly also with an ER-mitochondria contact site reporter (Figure 2C). In addition, Aim21 was identified as one of its HCPPs (Figure 1B, 1C, 2A), and since this protein is required for proper mitochondrial movement (Hess et al., 2009), we reasoned this may be at least part of the missing link between Lam5 and mitochondria. Indeed, we observed a unique colocalization between these proteins (Figure 2D). More work is required to understand the relationship between Lam5, Aim21, and ER-mitochondria contact sites, and we cannot rule out that other factors are at play. Vps35, a protein identified as an HCPP of both Lam5 and Lam6 may be one of these factors. Despite it being a highly-conserved component of the retromer complex in both yeast and humans, it is also known to regulate mitochondrial dynamics (Cutillo et al., 2020).One way it does this is via the removal of inactive human DNM1L (the ortholog of yeast Dnm1), which promotes mitochondrial fission (Wang et al., 2016). This would be interesting to explore in the future, especially in the functional context of Lam5 and Lam6.

To investigate the functionality of Lam5 at ER-mitochondria contacts, we used Seahorse technology to assay bioenergetics. Here, we found that the loss of Lam5 led to increased respiration when the cells were grown in glucose (Figure 2F), as opposed to Lam6, which appears to be required for maintaining normal respiration levels independent of the carbon source (Figure 2E, 2F). Evidence suggests these results are conserved, since loss of one of the human Lam5/6 orthologs, GRAMD1C, also leads to increased oxidative phosphorylation (Ng et al., 2022).

Interestingly, another Lam5/6 ortholog, GRAMD1B has been proposed to transfer cholesterol between the trans-Golgi network (TGN) and the ER (Naito et al., 2023). This is in line with localization data from yeast (Weill et al., 2018a) and our identification of Ret3, a COPI component, as a Lam5 HCPP. The role of Lam5 and Lam6 at multiple contacts is particularly intriguing, and, especially for Lam5, exactly how these different pools are maintained and regulated remains an open question. It does seem, however, that while there is some overlap in some of the locations and functions of the Lam5 and Lam6 paralogs, they each clearly have distinct characteristics. This is corroborated by our data (Figure 1B, 1C) and previous work on Lam6 interactors (Elbaz-Alon et al., 2015; Murley et al., 2015), which would indicate they do not interact with each other to form a complex. This is in contrast to the Lam1-4 family members, which appear to be in close proximity to each other (Figure 1B, 1C (Murley et al., 2017)).

Other HCPPs of Lam1-4 included several PM-resident proteins (Figure 3A), whose enrichment was greater for Lam2/4, relative to Lam1/3 (Figure 1B, 1C, 3A). This matched our microscopy observations, where Lam2/4 seem to be specifically localized to the peripheral ER, whereas Lam1/3 can also be seen at the perinuclear ER (Figure 3B). The different structural domains of Lam1/3, including the BAR domain, may explain these differences, however our proximity mapping did not yield any insights into the perinuclear Lam1/3 subdomain, and this would be of interest to resolve in the future.

The PM-resident Sso1/2 paralogs – which due to their high sequence similarity could not be differentiated from one another – were particularly curious HCPPs of Lam1-4. This is because in addition to being t-SNARE proteins for vesicle fusion, Sso1 and its human ortholog, STX1A, are also involved in *non-fusogenic* ER-PM tethering together with the ER-resident, Sec22 (yeast)/SEC22b (human) (Petkovic et al., 2014). In cells depleted for both SSO proteins, the localization of Lam1-4 was drastically reduced to a few puncta per cell (Figure 3C). Since Lam1-4 mediate retrograde sterol transfer from the PM to the ER, their perturbation renders cells sensitive to AmphotericinB (Gatta et al., 2015), an anti-fungal drug which affects PM sterols (Anderson et al., 2014). Since the deletion of both *sso1* and *sso2* is synthetic lethal, we could not assess AmphotericinB sensitivity in the double mutant strains (*GAL1pr-SSO1/ Δsso2)*. However, we could see that even the single mutant of *Δsso2* alone challenged by incubation at 37°C, grew slower on media containing the anti-fungal compared to controls (Supplementary figure 3B).

Collectively, our data suggest that the SSO proteins play an important role in regulating the ER-PM LAM contact sites. Exactly how this happens is likely to be a complex output of interactions between different protein domains and different lipid species, and this should be addressed in future experiments. Our hypothesis is that because human STX1A can cluster the anionic phosphorylated PI, PIP2, in the PM (Honigmann et al., 2013), then the SSO proteins may help promote Lam1-4 PM tethering through PIP2-PH domain interaction (Figure 4). Alternatively, Lam1-4 may physically associate with the SSO proteins, or there could even be a combination of protein and lipid-based interactions. In either case, PIP2 sequestration is important since its presence accelerates sterol transfer for yeast Lam2/4 and human GRAMD1B StART domains (Horenkamp et al., 2018; Jentsch et al., 2018). Furthermore, cholesterol can cluster STX1A (Murray and Tamm, 2009) and therefore it is possible that the lipid-binding requirements and properties of the SSOs may create an optimal and complementary environment for Lam1-4 association and lipid transfer.

More generally, by uncovering novel HCPPs for the LAM family we revealed both overlapping and specific properties for its paralogous members. We shed light on the localization and non-redundant function of the relatively uncharacterized protein, Lam5, and discovered the SSO proteins as novel, shared regulators of Lam1-4 ER-PM localization. Future work will tell if the SSO proteins are the direct tethering molecules for Lam1-4, however this discovery highlights the need to explore whether different contact site machineries can cooperate and coordinate with one another. With many different proteins and protein families mediating ER-PM tethering (Manford et al., 2012), we should question if, how, and under which conditions these machineries can work together. There is some evidence to suggest that tethering proteins collaborate in human cells, where GRAMD2 colocalizes with the ER-PM extended-synaptotagmin 2/3 tethers (Besprozvannaya et al., 2018). Lastly, several contact site proteins are known to be mutated in different human disorders. For example, a disease-linked mutation in GRAMD1B affects a residue conserved to yeast and is known to hinder anionic lipid detection by the protein’s PH-like domain (Ercan et al., 2021). These points emphasize the importance of elucidating the precise molecular mechanisms behind how these proteins work – for understanding both fundamental principles of cellular organization and potential implications for human health.

## Materials and Methods

### Yeast strains

The strains used in this work were either picked and verified from libraries, or constructed using the lithium acetate-based transformation protocol (Gietz and Woods, 2002). All strains are listed in Supplementary table 3, together with references.

### Yeast growth

Yeast cells were grown on solid media containing 2.2% agar or liquid media. YPD (2% peptone, 1% yeast extract, 2% glucose) was used for cell growth if only antibiotic selections were required, whereas synthetic minimal media (SD/Gal; 0.67% [w/v] yeast nitrogen base (YNB) without amino acids and with ammonium sulphate or 0.17% [w/v] YNB without amino acids and with monosodium glutamate, 2% [w/v] glucose (for SD) or 2% [w/v] galactose (for SGal), supplemented with required amino acid) was used for auxotrophic selection. Antibiotic concentrations were as follows: nourseothricin (NAT, Jena Bioscience) at 0.2g/l; G418 (Formedium) at 0.5g/l; and hygromycin (HYG, Formedium) at 0.5g/l. Yeast grown for transformation or protein extraction was first grown in liquid media with full selections overnight at 30°C and subsequently back-diluted into YPD/SD media to an OD_600_ of ∼ 0.2. Cells were collected after at least one division but before reaching an OD_600_ of 1 and either immediately used for transformation or snap-frozen for later processing.

For mass spectrometry, strains encoding TurboID-HA-LAM proteins on the background of ABOLISH were grown overnight at 30°C in SD liquid media supplemented with: an amino acid mix without leucine and histidine; G418; HYG; and 1mM auxin (Sigma, #13750). The following day, cells were back-diluted to an OD_600_ of ∼0.15 and collected by centrifugation after ∼5h (before reaching on OD_600_ of 1). Media for back-dilution was SD containing a complete amino acid mix, 1mM auxin and 100nM biotin (Supelco, #47868), as determined in (Fenech et al., 2023). Cell pellets were washed once in 1ml LC-MS/MS-grade H_2_O before being snap-frozen for later processing.

For imaging, the yeast strains were grown overnight in 100μl SD/SGal media (with appropriate amino acid and antibiotic selections) in round-bottomed 96-well plates at 30°C with shaking. For the non-SSO experiments, cells grown in SD-based media were back-diluted (5μl culture into 100μl fresh media) into SD (with complete amino acids) and grown for ∼4h at 30°C with shaking. For the SSO experiments, cells grown in either SD or SGal-based media were back-diluted into the same respective media and grown for another overnight. The following day, the cells were back-diluted into either SD or SGal (with complete amino acids), respectively, and grown for ∼4h. After the 4h growth period, cells were processed for imaging (see protocol below).

For the Seahorse assays, cells were initially grown for 24h on YPD agar plates supplemented with G418 at 30°C. Cells were then grown in liquid YPD (2% Glucose) or YPGal (1% Galactose) media overnight at 30°C with slight agitation. After 12h and on the day of the measurement, cells were back-diluted to an OD_600_ of 0.5 and grown for an additional 2.5h at 30°C with slight agitation, in either YPD or YPGal.

### Protein extraction and SDS-PAGE analysis

Protein extraction, sample preparation, SDS-PAGE electrophoresis, Western blotting and imaging were performed as described in (Eisenberg-Bord et al., 2021). Briefly, cell pellets were resuspended in 8M urea-based lysis buffer and subject to glass bead-beating. Lysates were denatured with SDS (final concentration ∼2%) and incubated at 45°C for 15min. Denatured lysates were centrifuged to separate cell debris. The resulting supernatants were reduced with sample buffer containing DTT (final concentration ∼25mM) and incubated at 45°C for 15min. Sample was separated on pre-cast 4-20% gradient gels (Bio-Rad) which were transferred onto nitrocellulose membrane using the Trans-Blot Turbo transfer system (Bio-Rad). Membranes were blocked in SEA BLOCK buffer (Thermo Scientific), incubated overnight at 4°C with primary antibodies (anti-HA, 1:1000, BioLegend, #901502; anti-myc, 1:3000, Abcam, #ab9106; anti-Histone H3, 1:5000, Abcam, #ab1791), washed, and finally incubated with fluorescent secondary antibodies (goat anti-rabbit IgG 800, 1:10000,Abcam, #ab216773; goat anti-mouse IgG 680, 1:10000, Abcam, #ab216776) for 1h at room temperature. After washing, the probed membranes were imaged on the LI-COR Odyssey Infrared Scanner.

### Affinity purification and LC-MS/MS sample preparation for TurboID-HA-tagged protein samples

Samples were prepared exactly as reported in (Fenech et al., 2023). Briefly, cell pellets were resuspended in 400μl lysis buffer (150mM NaCl, 50mM Tris–HCl pH 8.0, 5% Glycerol, 1% digitonin (Sigma, #D141), 1mM MgCl_2_, 1 x protease inhibitors (Merck), benzonase (Sigma, #E1014)) and lysed by 6 × 1min maximum speed cycles on a FastPrep-24™ cell homogenizer (MP Biomedicals) using 1mm silica beads (lysing matrix C, MP Biomedicals). Lysates were cleared by centrifugation and subject to affinity purification overnight at 4°C with streptavidin-conjugated magnetic beads (Cytiva, #28985799). The beads were subsequently washed twice in 2% SDS, twice in 0.1% SDS, and lastly, twice in basic wash buffer (150mM NaCl, 50mM Tris–HCl pH 8.0). Elution buffer (2M urea, 20mM Tris–HCl pH 8.0, 2mM DTT and 0.25μg/μl trypsin/sample) was added to the beads, followed by alkylation, and digestion overnight. The following morning 0.25μg/μl trypsin was added to each sample and incubated for a further 4h. Peptides were acidified and desalted using Oasis HLB, μElution format (Waters, Milford, MA, USA). The samples were vacuum dried and stored at −80°C until further analysis.

### LC-MS/MS settings, analysis and raw data processing (for TurboID-HA-tagged protein samples)

Settings, analysis and data processing were carried out exactly as described in (Fenech et al., 2023). In short, samples were run on a Q Exactive HF instrument (Thermo Scientific) and data were acquired in data-dependent acquisition (DDA) mode. Raw data were processed using MaxQuant v1.6.6.0 and searched with the Andromeda search engine against the SwissProt *S. cerevisiae* database (November 2018, 6049 entries). The generated LFQ intensities were used for subsequent analysis on Perseus v1.6.2.3 and Student’s *t*-tests were carried out between appropriate groups to identify significantly enriched proteins. The raw datasets were deposited on the ProteomeXchange Consortium via the PRIDE partner repository (Perez-Riverol et al., 2021), under the identifier PXD051047. The LAM HCPPs are listed in Supplementary table 1, together with information on their localization from (Dubreuil et al., 2018).

### Generation and culture of GRAMD1A^FLAG^ cell lines

Constructs enabling expression of GRAMD1A with a C-terminal FLAG tag were generated by amplification of gene-specific PCR fragments using HEK293T cDNA. Primers were designed based on the NCBI (National Center for Biotechnology Information) sequence (NM_020895.5). Amplicons and pcDNA5/FRT/TO (Thermo Fisher Scientific) were digested with appropriate enzymes, ligated and the final constructs confirmed by sequencing. Human embryonic kidney (HEK) cell lines that inducibly express GRAMD1A^FLAG^ using the HEK293T-Flp-In™ T-Rex™ system were generated according to manufacturer’s instructions (Thermo Fisher Scientific). Single clones were selected and confirmed. Expression of the construct was induced using 1μg/ml doxycycline (Sigma-Aldrich, D9881) at 14h prior to harvest. For interactomic experiments, a stable-isotope labeling of amino acids in cell culture (SILAC) approach was taken whereby cells were grown for five passages in DMEM medium lacking arginine and lysine, supplemented with 10% (v/v) dialyzed fetal bovine serum, 600mg/l proline, 42mg/l arginine hydrochloride (or ^13^C_6_, ^15^N_4_-arginine in ‘heavy’ media) (Cambridge Isotope Laboratories) and 146mg/l lysine hydrochloride (or ^13^C_6_, ^15^N_2_-lysine in ‘heavy’ media) (Cambridge Isotope Laboratories).

### Crosslinking, immunoprecipitation and LC-MS/MS sample preparation of GRAMD1A^FLAG^

To stabilize any potential GRAMD1A interactions, cells were washed twice in PBS with 0.1mM CaCl_2_ and 1mM MgCl_2_ (PBS++) and then crosslinked with 0.5mM ethylene glycol bis(succinimidyl succinate) (EGS) in PBS++ for 30 min at 37°C. Cells were then washed with PBS++ and the reaction was quenched using 10mM NH_4_Cl in PBS++ for 10 min at 37°C. Cells were harvested, washed in PBS and resuspended in solubilization buffer (50mM Tris–HCl pH 7.4, 150mM NaCl, 10% (v/v) glycerol, 1mM EDTA, 1% (v/v) digitonin, 1mM PMSF, 1 x protease inhibitors (Roche)) at a protein/buffer ratio of 2mg/ml. Cells were solubilized for 30min at 4°C with mild agitation. Unsolubilized cells were sedimented by centrifugation at 12,000g for 15min at 4°C. Solubilized protein was added to pre-equilibrated anti-FLAG agarose affinity resin (Sigma, A2220) and incubated for 90min at 4°C on a rotating wheel. Affinity resins were washed extensively with buffer containing 0.3% (v/v) digitonin. Bound proteins were eluted with 5μg/ml FLAG peptide (Sigma, F3290) in buffer and subsequently separated on 4-12% NuPAGE Novex Bis-Tris Minigels (Invitrogen). Gels were stained with Coomassie Blue for visualization purposes, and each lane sliced into 21 equidistant slices regardless of staining. After washing, gel slices were reduced with dithiothreitol (DTT), alkylated with 2-iodoacetamide and digested with Endopeptidase Trypsin (sequencing grade, Promega) overnight. The resulting peptide mixtures were then extracted, dried in a SpeedVac, reconstituted in 2% acetonitrile/0.1% formic acid (v/v) and prepared for nanoLC-MS/MS as described previously (Atanassov and Urlaub, 2013).

### LC-MS/MS settings, analysis and raw data processing (for GRAMD1A^FLAG^ samples)

For mass spectrometric analysis, samples were enriched on a self-packed reversed phase-C18 precolumn (0.15mm ID x 20mm, Reprosil-Pur120 C18-AQ 5µm, Dr. Maisch) and separated on an analytical reversed phase-C18 column (0.075mm ID x 200mm, Reprosil-Pur 120 C18-AQ, 3µm, Dr. Maisch) using a 30min linear gradient of 5-35 % acetonitrile/0.1% formic acid (v:v) at 300nl/min). The eluent was analyzed on a Q Exactive hybrid quadrupole/orbitrap mass spectrometer (ThermoFisher Scientific) equipped with a FlexIon nanoSpray source and operated under Excalibur 2.4 software using a data-dependent acquisition (DDA) method. Each experimental cycle was of the following form: one full MS scan across the 350-1600 m/z range was acquired at a resolution setting of 70,000FWHM, and AGC target of 1x10^6^ and a maximum fill time of 60ms. Up to the 12 most abundant peptide precursors of charge states 2 to 5 above a 2x10^4^ intensity threshold were then sequentially isolated at 2.0FWHM isolation width, fragmented with nitrogen at a normalized collision energy setting of 25%, and the resulting product ion spectra recorded at a resolution setting of 17,500FWHM, and AGC target of 2x10^5^ and a maximum fill time of 60ms. Selected precursor m/z values were then excluded for the following 15s. All gel fractions were acquired with two technical injection replicates.

Raw MS files were processed using MaxQuant (version 1.5.7.4) and MS/MS spectra were searched against the UniProtKB database human reference proteome (February 2017) using default SILAC settings for K8/R10 heavy channels, and the ‘Requantify’ option enabled for improved quantitation. Statistical analysis was performed in Perseus software (version 1.6.15.0); the Significance B test was used to establish significant abundance changes on the protein group level. The raw datasets were deposited on the ProteomeXchange Consortium via the PRIDE partner repository (Perez-Riverol et al., 2021), under the identifier PXD051014. Significantly enriched/depleted proteins are listed in Supplementary table 2.

### Imaging and organelle staining

After back-dilution of strains for imaging (see *Yeast Growth* methods above), 50μl of cell culture was transferred to a 384-well glass-bottomed plate (Azenta Life Sciences) coated with Concanavalin A (ConA, Sigma, 0.25mg/ml) and incubated at RT for ∼20min. Cells were then washed twice in SD (or SGal for Figure 3C and 3D) supplemented with a complete amino acid mix and imaged in the same media. If MitoView™405 (Biotium, #70070) / FM™4-64 (Invitrogen, #T13320) dyes were being used for mitochondrial/vacuolar staining, then prior to washing, the media was removed and 50μl of SD supplemented with a complete amino acid mix containing the required dye/s was added and incubated at RT for 15min. Dye concentrations were: 500nM; 50nM; and 16μM, respectively.

For Figure 2C and 2D, images were obtained using a VisiScope Confocal Cell Explorer system composed of a Yokogawa spinning disk scanning unit (CSU-W1) coupled with an inverted Olympus IX83 microscope. Single focal plane images were acquired with a 60x oil lens and were captured using a PCO-Edge sCMOS camera, controlled by VisiView software (V3.2.0, Visitron Systems; GFP/Venus at 488 nm and mCherry at 561 nm). Images were transferred to ImageJ (Schindelin et al., 2012) for slight contrast and brightness adjustments. The contrast and brightness settings for the ER-mitochondria CS reporter images (Figure 2C) and for Aim21-mScarlet images (Figure 2D) were set relative to the mCherry-/GFP-Lam5 strains to enable direct comparison with the mCherry-/GFP-Lam6 strains.

For all other microscopy-based figures, images were obtained using an automated inverted fluorescence microscope system (Olympus) containing a spinning disk high-resolution module (Yokogawa CSU-W1 SoRa confocal scanner with double micro lenses and 50 μm pinholes). Several planes were recorded using a 60x oil lens (magnification 3.2x, NA 1.42) and with a Hamamatsu ORCA-Flash 4.0 camera. Fluorophores were excited by a laser and images were recorded in three channels: GFP (excitation wavelength 488 nm, emission filter 525/50 nm), mCherry / mScarlet / FM™4-64 (excitation wavelength 561 nm, emission filter 617/73 nm) and MitoView™405 (excitation wavelength 405 nm, emission filter 447/60). Image acquisition was performed using scanR Olympus soft imaging solutions version 3.2. Images were transferred to ImageJ (Schindelin et al., 2012), for slight contrast and brightness adjustments to each individual panel.

### Split-Venus assay

Generation and imaging of strains expressing *TEF2pr*-mCherry-tagged Lam5 or Lam6 together with an ER-mitochondria contact site reporter (Pho88-VC in the ER membrane and Tom70-VN in the outer mitochondrial membrane) was carried out exactly as detailed in (Castro et al., 2022). Briefly, a strain expressing Pho88-VC and Tom70-VN was crossed against strains picked from the *TEF2pr*-mCherry library (Weill et al., 2018b; Yofe et al., 2016) and then sporulated and selected for haploids containing both traits using automated methods (Cohen and Schuldiner, 2011; Tong and Boone, 2006). The resulting haploid strains were imaged on a Yokogawa spinning disk scanning unit (see under *‘Imaging and organelle staining’*).

### Seahorse metabolic analysis

One day prior to measurement, a Seahorse XFe96/XF Pro Cell Culture microplate (Agilent) was coated with 0.1mg/ml Poly-D-Lysine (Sigma Aldrich) and incubated at 4°C overnight. Seahorse XF Calibrant (Agilent) was added to the Seahorse XFe96/XF Pro Sensor Cartridge plate (Agilent) and incubated overnight in a non-CO_2_ incubator at 37°C. The Seahorse XFe96 Analyzer (Agilent) was set to 30°C. On the day of measurement, cells were pelleted by centrifugation at 500g for 5min to remove growth media, after which they were resuspended in assay medium (0.67% yeast nitrogen base, 2% potassium acetate, and 2% ethanol) as described in (Zhang et al., 2022), to an OD_600_ of 0.1. 180μl of cell suspension per well was added to the Seahorse XFe96/XF Pro Cell Culture microplate followed by incubation at 30°C for 30min. Measurements were taken under basal conditions (from 0min) and upon the addition of 40mM CCCP (at 18min; Sigma, #C259), and 2.5mM Antimycin A (at 36min; Sigma #A8674) in the Seahorse Analyzer. Three 6min cycles of mixing (for 3min) and measuring (for 3min) time were allotted to each condition. Data analysis was done using Graph Pad Prism (V10.2.0) and the data presented is the average of six independent experiments. Graphs are plotted showing the standard error of the mean (SEM). Statistical significance was determined using two-way ANOVA with Tukey’s multiple comparison. *P*-values ≤ 0.05, ≤ 0.01, ≤ 0.001, or ≤ 0.0001 are shown as *, **, ***, or ****, respectively. *P*-values >0.05 are not significant (ns).

### AmphotericinB serial dilution growth assay

Cells were grown overnight in liquid YPD media supplemented with NAT. The following day, cells were back-diluted in YPD to an OD_600_ of 0.2 and grown for ∼3h at 30°C. Serial dilutions were prepared exactly as described in (Castro et al., 2022) and cells were plated on SD + complete amino acid agar plates with or without 1μg/ml AmphotericinB (Sigma, #A2932). Cells were incubated for two days at 37°C and then imaged.

## Data availability

Yeast TurboID-HA-tagged proximity labeling LC-MS/MS data is available via ProteomeXchange with identifier PXD051047. Human GRAMD1A-FLAG IP-LC-MS/MS data is available via ProteomeXchange with identifier PXD051014.

## Acknowledgements

We thank Dr. Ehud Sass, Sivan Arad and Noga Preminger from the Schuldiner lab for important feedback on this manuscript, and Reut Ester Avraham for making all the yeast media. This project was supported by the Deutsche Forschungsgemeinschaft through the SFB1190 (P11, MS; P13, PR; Z02, CL). Emma Fenech was supported by a senior postdoctoral award from the Weizmann Institute of Science. Maya Schuldiner is an incumbent of Dr. Gilbert Omenn and Martha Darling Professorial Chair in Molecular Genetics.

## Author contributions

Emma J Fenech conceptualized the project, designed and performed experiments, analyzed data, and wrote the original manuscript draft. Meital Kupervaser ran TurboID-HA-LAM samples and processed the raw LC-MS/MS data. Angela Boshnakovska performed and analyzed Seahorse experiments. Shani Ravid performed microscopy and drop assays. Inês Gomez Castro performed the split-Venus assay. Yeynit Asraf facilitated enhanced-resolution microscopy. Sylvie Callegari performed GRAMD1A-FLAG pulldowns and the LC-MS/MS was performed by Christof Lenz. Peter Rehling supervised the project as well as acquired resources and funding. Maya Schuldiner conceptualized and supervised the project, acquired resources and funding, and wrote the original manuscript draft. All authors contributed to reviewing and editing the manuscript.

---

**Supplementary figure 1:**
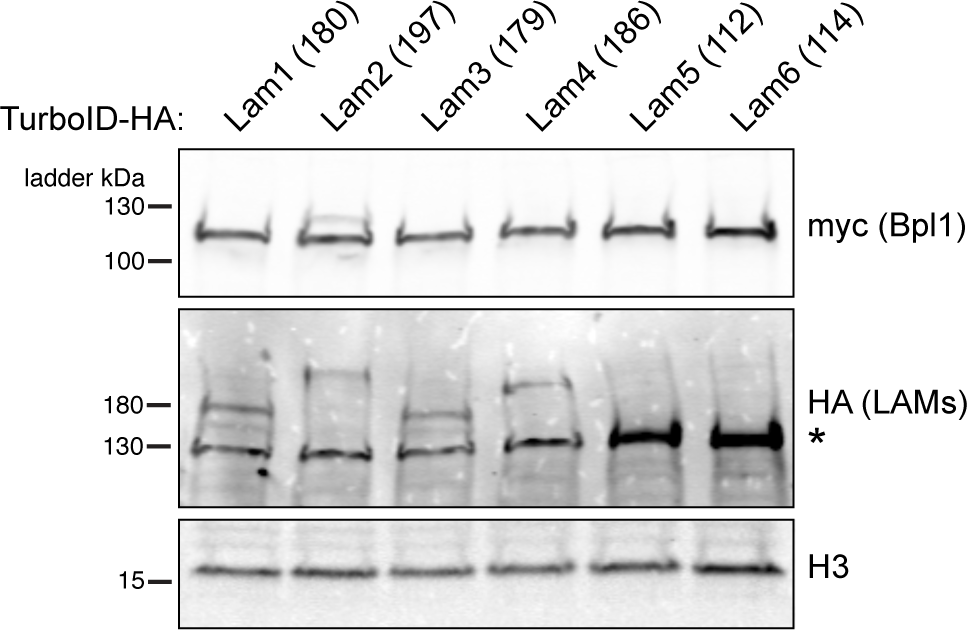
Western blot analysis confirming expression of TurboID-HA-tagged LAM proteins on the ABOLISH background. The anti-myc blot shows expression of Bpl1 tagged on its C-terminus with AID*-myc. Anti-histone (H3) was used as a loading control. The predicted molecular weights for the tagged LAM proteins are indicated in parentheses. A prominent non-specific band (marked by an asterisk) is at the same molecular weight as tagged Lam5 and Lam6, but both can be seen above this background as a much more intense signal.

**Supplementary figure 2:**
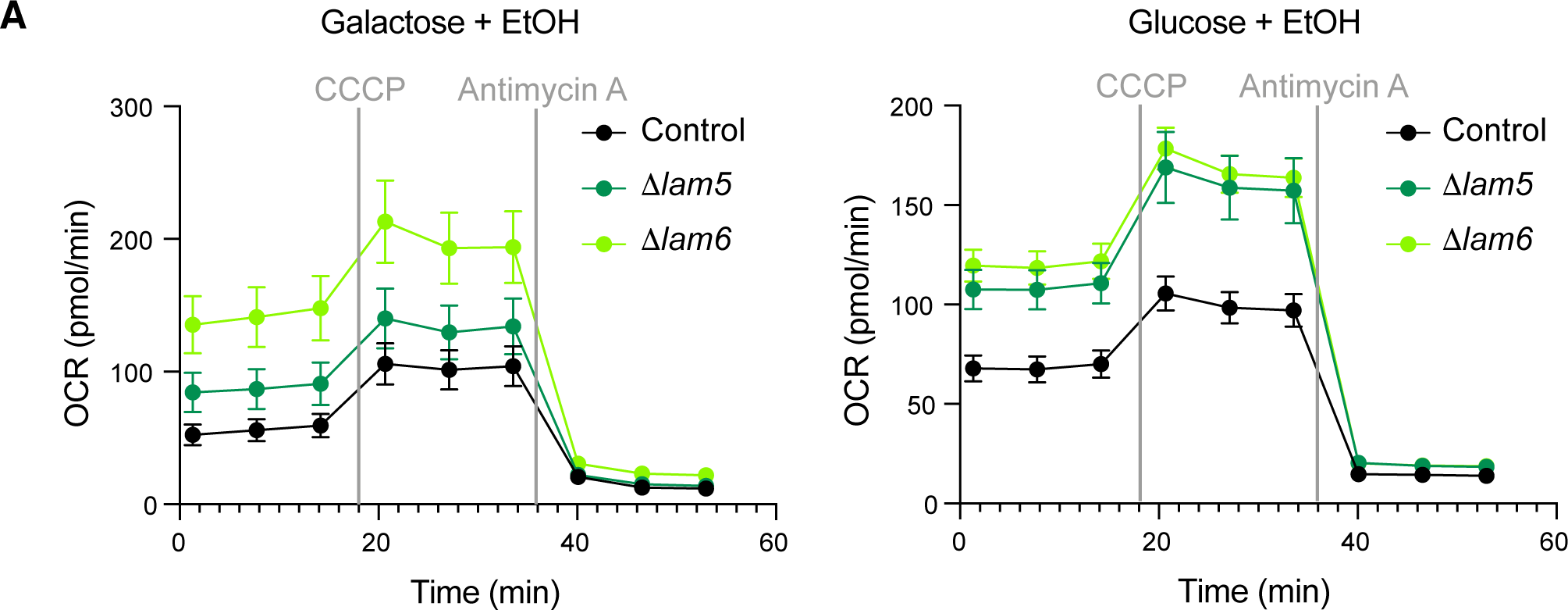
(A) Graphs showing the OCR profiles from 0 to 53 minutes (min) of control (black), *Δlam5* (dark green), and *Δlam6* (light green) strains grown in either galactose (left) or glucose (right) mixed with ethanol. The grey vertical lines indicate the points at which CCCP (a mitochondrial uncoupler which induces maximal OCR) and Antimycin A (a complex III inhibitor which allows background OCR to be measured) were added. The three time points taken prior to CCCP addition are considered ‘basal’, whereas the three time points after CCCP addition are ‘maximal’ (Fig 2E and F). Error bars show the SEM from all six biological replicates.

**Supplementary figure 3:**
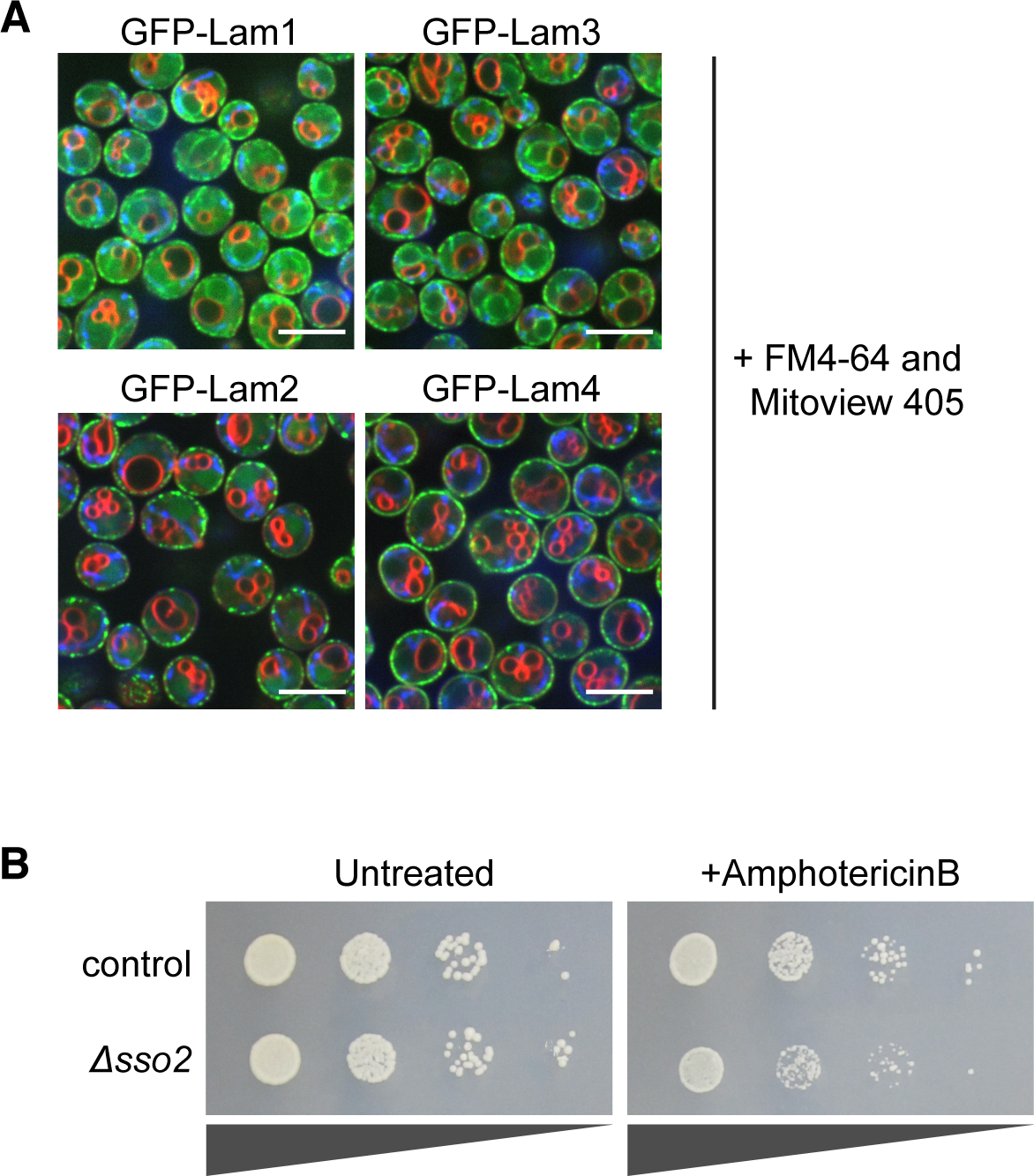
(A) Images from main figure 3A including vacuolar (FM™4-64) and mitochondrial (MitoView™405) dyes. Unlike Lam5 and Lam6, virtually no colocalization between these LAM proteins and the dyed organelles was observed. Scale bars are 5μm. (B) Drop assay of control and *Δsso2* strains grown with or without AmphotericinB at 37°C. Loss of *SSO2* affects growth in the presence of AmphotericinB.

